# Developing a safe 15-hour preservation protocol of donor kidneys using normothermic machine perfusion

**DOI:** 10.1101/2023.09.12.555854

**Authors:** John P Stone, William R Cowey, Corban JT Bowers, Amy F Stewart, Erin R Armstrong, Marc Clancy, Timothy R Entwistle, Jorge del Pozo, Kavit Amin, James E Fildes

## Abstract

**Introduction:** *Background:* Normothermic machine perfusion (NMP) offers a superior alternative to existing hypothermic preservation strategies but is currently limited to 1-3 hours. Extending the time a kidney can be sustained using this technology could electivise transplantation, and enable physiological assessments of renal function. We aimed to develop a protocol that allows the safe preservation of donor kidneys for 12 hours using this technique.

*Methods:* Porcine kidneys (n=20) were retrieved and flushed with 1L preservation solution before being stored on ice. Following a cold ischaemic time of 3.5 hours, kidneys were placed onto a NMP circuit and perfused for 12 hours. Renal haemodynamics, biochemistry and urine output were recorded and analysed. At the end of perfusion, kidneys were scored based on the clinical assessment score and their suitability for transplant determined. Biopsies were collected at the end for histological assessment.

*Results:* All kidneys were successfully perfused with immediate recordable renal blood flow (RBF). RBF continually improved over the course of the perfusions, peaking at 12 hours, and negatively correlated with intra-renal resistance. Perfusate sodium concentrations remained stable and within physiological parameters. Sodium bicarbonate increased over time with a corresponding decrease in lactate concentrations, demonstrating active renal gluconeogenesis and Cori cycle processes. Urine production began immediately in all kidneys and was sustained throughout, indicating active renal function. Under the clinical perfusion assessment score, all kidneys received a score of 1 and would be considered suitable for transplantation. Histological assessment revealed kidneys were injury free with REMUZZI scores of 0 in all samples.

*Conclusion:* We have developed an NMP protocol that safely preserves donor kidneys for over 15 hours. Successful perfusion was achieved with stable haemodynamics, blood-perfusate biochemistry, and maintained urine output. Importantly, kidneys remained in optimal health, with no evidence of injury. This protocol may enable the electivisation of transplantation, while reducing ischaemic injury associated with static cold storage.

## Introduction

Kidney transplantation is the gold standard of care for all patients with end-stage renal disease. In the UK there were 2,868 single kidney donations in 2022; this includes live donation, donation after cardiac death (DCD) and donation after brain death (DBD) (1). However, at this time there were 4,997 patients on the kidney transplant waiting list, meaning 2,129 transplantable patients had to remain on other treatment options, often including dialysis. Such treatments carry a significant socioeconomic burden, increasing demands on healthcare providers, reducing the quality of life for the patient and reducing life expectancy significantly (2,3).

The primary limitation to successful kidney transplantation is the lack of suitable organs, forcing the assessment and use of older and more comorbid donors, the kidneys from which appear far more vulnerable to the multiple injurious processes associated with storage and transplantation. These secondary complications include ischaemia reperfusion injury (IRI), which if severe, manifests clinically as delayed graft function (DGF) or primary non-function post-transplant. Severe DGF is strongly associated with acute allograft rejection, but all recipients require life-long immunosuppression, which inherently increases the susceptibility to cancer and infection. It is well documented that the longer a donor kidney is exposed to static cold storage (SCS), which remains the current gold standard of preservation, the more severe the IRI is, in turn increasing the likelihood of DGF post-transplant (2). DGF alone increases the overall cost of transplantation, requiring extended hospital stays and renal support such as temporary dialysis, whilst also reducing graft survival.

Normothermic machine perfusion (NMP) is a technique that offers a more physiological approach to donor kidney preservation. Organs are placed into a chamber and a normothermic (37°C) blood-based perfusate is pumped across the vasculature at physiological pressures, restoring function and metabolism outside of the body. An extracorporeal membrane oxygenator is used to deliver oxygen to the perfusate and remove CO_2_. A pump (levitating centrifugal pump or rotary pump) is used to control arterial flow rate through the kidney. Currently, this technique is used clinically following a period of static cold storage (SCS) or machine cold perfusion to allow a short-term (1-3 hours) evaluation of kidney function, before making a decision on the suitability of an organ for transplantation. Donor kidneys that have been exposed to NMP for this short time have improved function, compared to donor kidneys that have been preserved solely via SCS (3,4). Once optimised, this technology has the potential to extend donor kidney preservation without ischaemia, cellular injury, or donor derived inflammation, potentially changing kidney transplant from an emergency procedure to elective surgery.

The aim of this study was to evaluate if a novel NMP protocol can safely perfuse donor kidneys for 12 hours without loss of physiological perfusion parameters. This was selected by the clinical team as a critical time threshold for donor kidney transplantation.

## 2 Methods

### Retrieval

Paired kidneys were retrieved from n=20 landrace pigs (Sus scrofa domesticus) as previously described (5). The pigs had a mean dry weight of 80kg and were all veterinary inspected and culled in accordance with the European union council regulation (EC) No 1099/2009 on the protection of animals at the time of killing. Briefly, pigs were electrically stunned followed by exsanguination. Approximately 5 litres of autologous blood was collected into a sterile container containing saline (Baxter Healthcare Ltd, UK), unfractionated heparin (Kent Pharma, UK), and antibiotics. A midline incision was made, and bilateral kidneys were inspected for infection and cysts, before being removed en-bloc and immediately placed on ice. Once the kidneys were placed on ice the renal artery was isolated and cannulated, and kidneys were then flushed with cold custodiol (Dr Franz Köhler Chemie Gmbh, Germany) solution supplemented with heparin at a pressure of approximately 80mmHg using a pressure bag. The time was recorded to mark the start of cold ischaemia. Whilst being flushed residual fatty tissue was removed from the kidney and the ureters were isolated and cannulated. At the end of the flush, kidneys were placed into a bag containing cold custodiol, and then stored in an ice slurry for 3 hours (SCS of 3 hours was selected as a clinically relevant time to transport a donor kidney to the recipient hospital prior to NMP). This represented a total time of 3.5hrs (including 30 minutes of back table preparation and SCS).

### Perfusion

Autologous blood was cell saved using a Cell Saver 5+ (Haemonetics,USA), and leukocyte depleted using RS1 filters (Haemonetics, USA). The RBCs were then added to a mix of drugs to make up the perfusate mix. The perfusate mix was then added to a pre-built circuit comprising of a bespoke kidney chamber, a reservoir and extracorporeal membrane oxygenator (Trilly Pediatrics, Eurosets, Italy), and a centrifugal pump (Eurosets, Italy). Additional attachments to the circuit included a heater (Eurosets, Italy) which was set to a target perfusate temperature of 39°C, flow probe, bubble sensor, temperature probe, pressure line, an oxygen line which was connected to a carbogen gas cannister (95% oxygen, 5% carbon dioxide) and syringe pumps. Once the perfusate mix was added to the circuit, the system was primed to ensure it was completely deaired. After the system was primed, multiple infusions were set up and started immediately prior to the kidney being connected.

Once the circuit was primed an arterial blood gas (GEM Premier 4000, Instrumentation Laboratory, USA) was performed to ensure parameters were within physiological range. Once achieved, kidneys were removed from cold storage and flushed with cold ringers (Baxter Healthcare Ltd, UK) to remove residual preservation solution. Kidneys were weighed before being attached to the circuit via the renal artery, marking the end of cold ischaemia. During perfusion, serial hourly arterial blood gases were recorded until the end of the experiment. Any deviation from reference physiological blood parameters (generated in-house) were corrected. Urine output (UO) was continually monitored and replaced with Ringer’s solution. At the end of the perfusion, kidneys were weighed and dissected using a sagittal cut to allow for macroscopic inspection.

### Clinical Assessment Score

Kidneys were scored according to the clinical assessment score validated by Hosgood *et al*. (2015) to determine suitability for transplantation (3). The criteria scores kidneys on a scale of 1-5 based on renal blood flow (RBF), UO and macroscopic appearance, where 1 is the highest quality kidney and 5 is the poorest. Macroscopic appearance is graded whereby grade I is a global pink appearance, grade II is a patchy appearance and grade III is a global mottled and purple/black appearance (scored 1, 2 and 3 respectively). A UO of less than 43ml and a mean RBF of less than 50ml/mg/100g both add 1 to the score.

### Histology

One needle biopsy from five randomly selected kidneys and one wedge biopsy from a further five randomly selected kidneys were sent to an external histology lab (Easter Bush Pathology, Edinburgh). Samples were processed and stained with haematoxylin and eosin (H&E). Histological evaluation was performed by an external experienced veterinary histopathologist. All stains were assessed qualitatively for signs of damage and tissue integrity. Kidneys were also scored according to Remuzzi score which assesses the severity of tissue damage based on presence of glomerular global sclerosis, tubular atrophy, interstitial fibrosis and arterial/arteriolar narrowing (6). Each factor is given a score of 0-3 where 0 is no evidence of and 3 is high severity. This gives a possible total histology score of between 0 and 12. A kidney with a score of 3 or less (mild) is considered adequate for single transplant, a score of 4 to 6 (moderate) for double transplant, and a score greater than 7 (severe) discarded. The five core needle biopsies were also gram stained and assessed for any signs of bacterial infection.

### Statistical Analysis

All numerical data were analysed and graphed using GraphPad Prism v.9.5.1. Data is presented as mean ± standard deviation. Data normality was determined by assessing mean, standard deviation, skewness and kurtosis and formal evaluation was performed using the Shapiro-Wilk test. To assess changes in variables over-time a One-way ANOVA or Mixed-effects analysis was used. The Pearson’s correlation coefficient was used to assess correlations between two variables. A Wilcoxon matched-pairs signed rank test was used to assess individual time-points. Data was considered significantly different if a p-value of ≤0.05 was observed.

## 3 Results

### Renal Haemodynamics

RBF was recordable in all kidneys within the first 15 minutes of perfusion, significantly increasing by 60 minutes (p=0.0074). Over the 12 hours, blood flow continually increased (p<0.0001), reaching peak RBF at 12 hours (85.7mls/min/100g ± 13.8, Figure 1A). Conversely, intra-renal resistance (IRR) continually reduced (p=0.0010, Figure 1B) and as expected, had a significant negative correlation with RBF (r=-0.955, p<0.0001). Mean arterial pressure within the renal artery was maintained at 82.5mmHg ± 0.49 (p=0.16, Figure 1C), demonstrating preserved autoregulation of RBF and vascular resistance ex-vivo.

**Figure 1.**
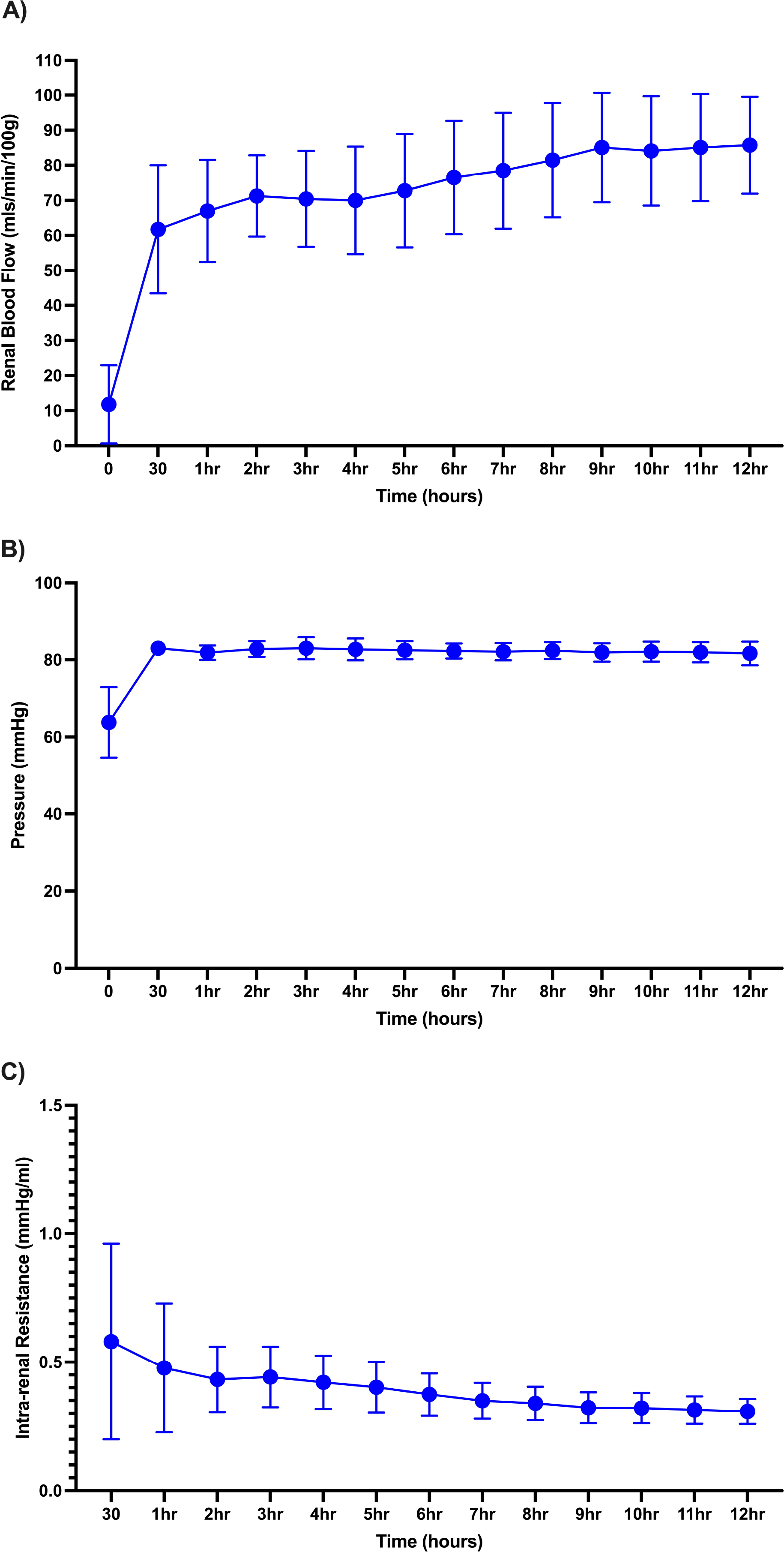
Renal hemodynamics remain stable during perfusion. Renal blood flow (A) continually increased whilst Intra-renal resistance declined (B). Pressure was maintained within physiological parameters (C).

### Renal Biochemistry

Overall, perfusate biochemistry remained stable and within physiological parameters (Figure 2A-E). Sodium concentrations within the perfusate were maintained at an average of 146.8 ± 3.48 with minimal variation across the 12 hours (p=0.047). In all perfusions, sodium bicarbonate was supplemented at baseline to achieve a pH within physiological ranges (7.4-7.53). This became slightly alkalotic during the perfusions (7.55 ± 0.06), likely as a result of increasing sodium bicarbonate concentrations (p<0.0001) as pCO_2_ remained within acceptable ranges (40.97 ± 7.73). Lactate levels were high in the immediate perfusion period owing to Ringer’s lactate being used as the base of the perfusion solution (10.86 ± 1.35). Once the kidneys were attached to the NMP circuit there was a significant reduction in lactate concentrations (p=0.0003). A significant correlation between diminishing lactate concentrations and the increase in sodium bicarbonate was observed, indicating active renal gluconeogenesis and Cori cycle function (r=-0.99, p<0.0001, Figure 3). Glucose was actively consumed during perfusion, requiring occasional supplementation to maintain euglycaemia. However, given the kidney is involved with gluconeogenesis, glycolysis, glucose filtration, and glucose reabsorption, glucose consumption could not be assessed.

**Figure 2.**
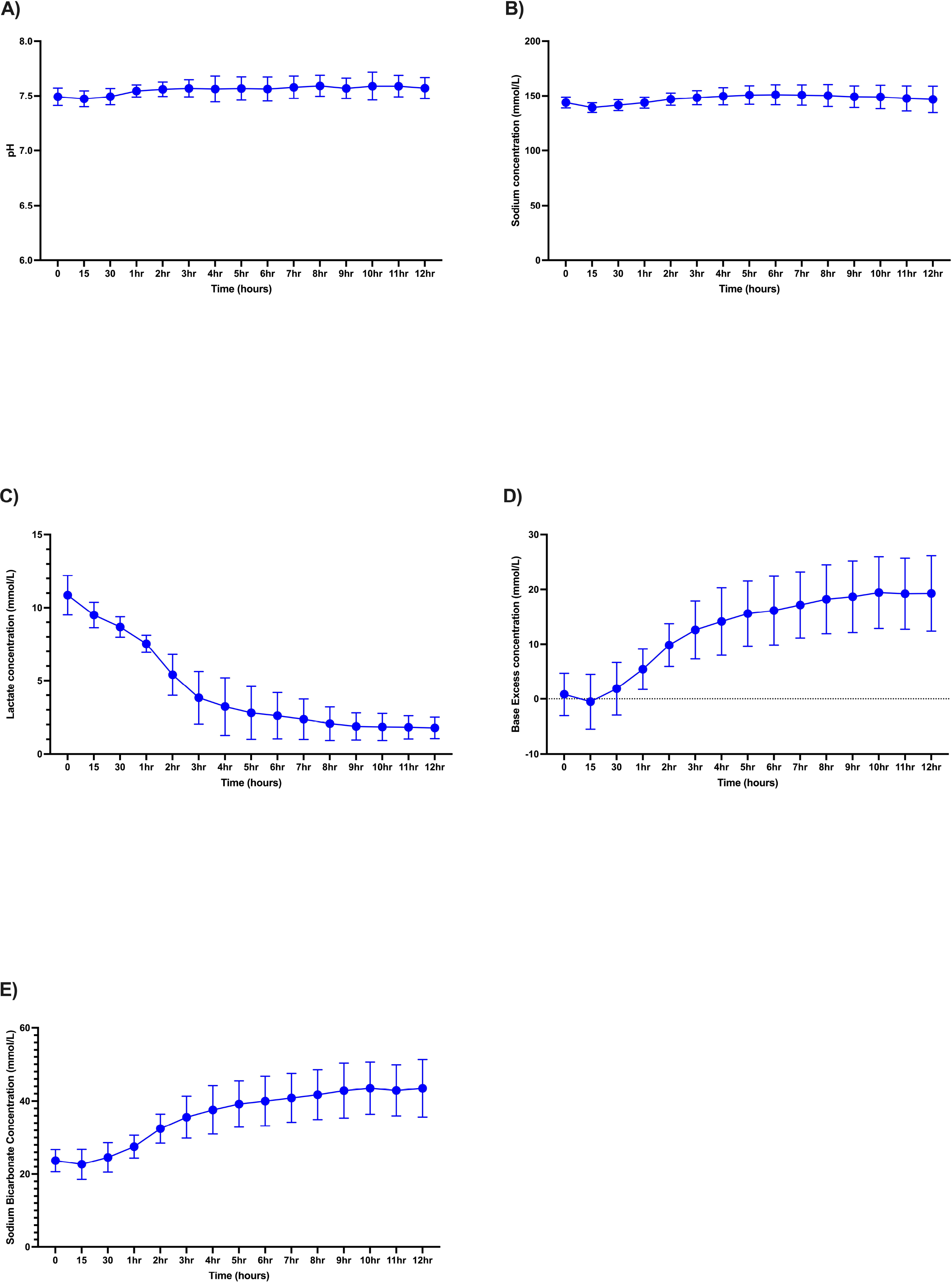
Kidneys remain within physiological biochemical parameters. pH and sodium concentrations were stable during 12 hours of perfusion (A,B). Lactate declined during perfusion (C), whilst base and sodium bicarbonate concentrations increased (D,E).

**Figure 3.**
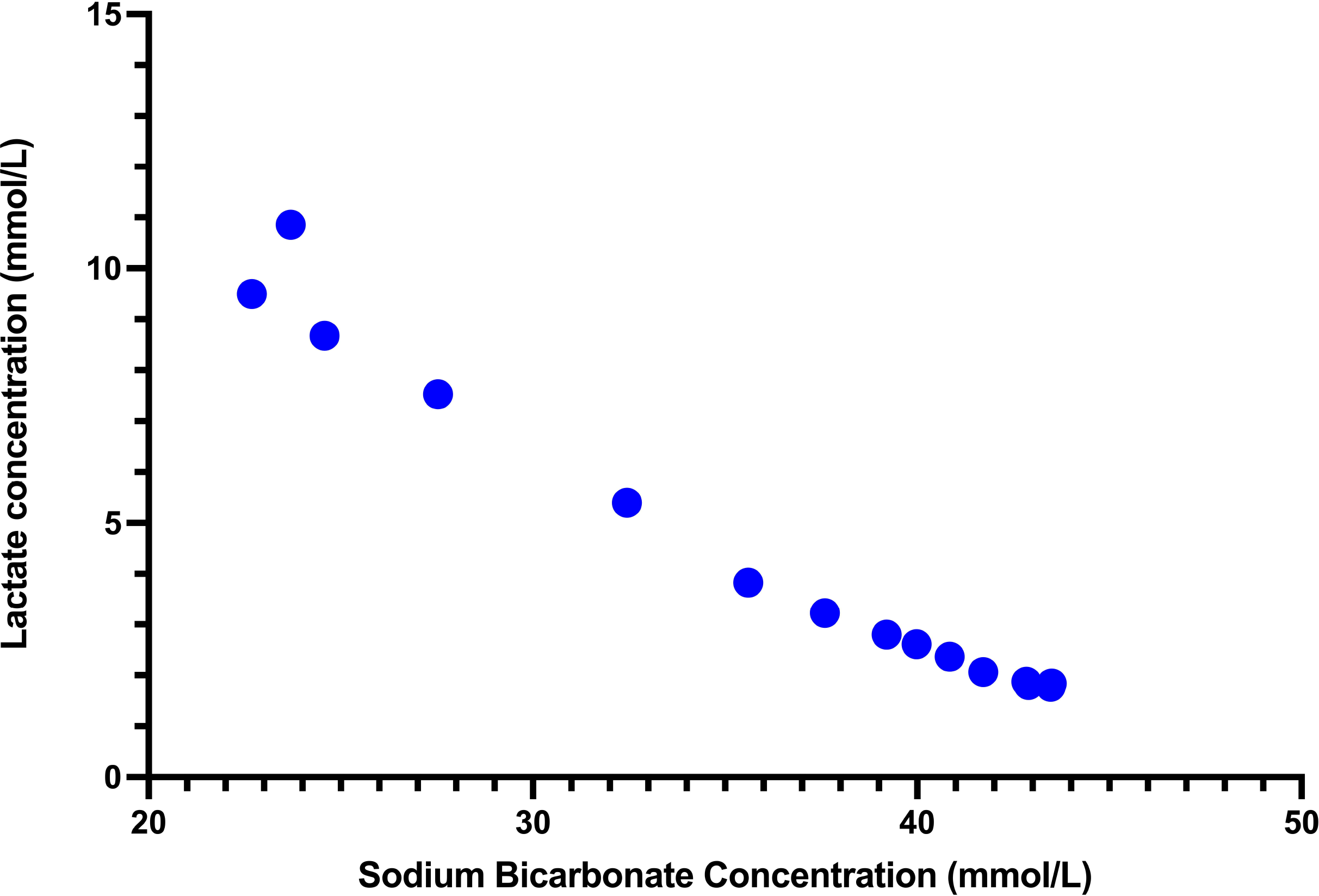
Sodium bicarbonate and lactate concentrations correlate over-time. A negative correlation is observed between sodium bicarbonate and lactate concentrations, with sodium bicarbonate increasing as lactate decreases.

### Renal Function

Urine production began in all kidneys immediately upon connection to the circuit (Figure 4). Urine production was highest in the first hour of perfusion with an average of 46.2mls **±** 54.1. This was maintained for the duration of the experiments with a mean total UO of 235.4mls ± 228.2, demonstrating restoration and sustained functionality.

**Figure 4.**
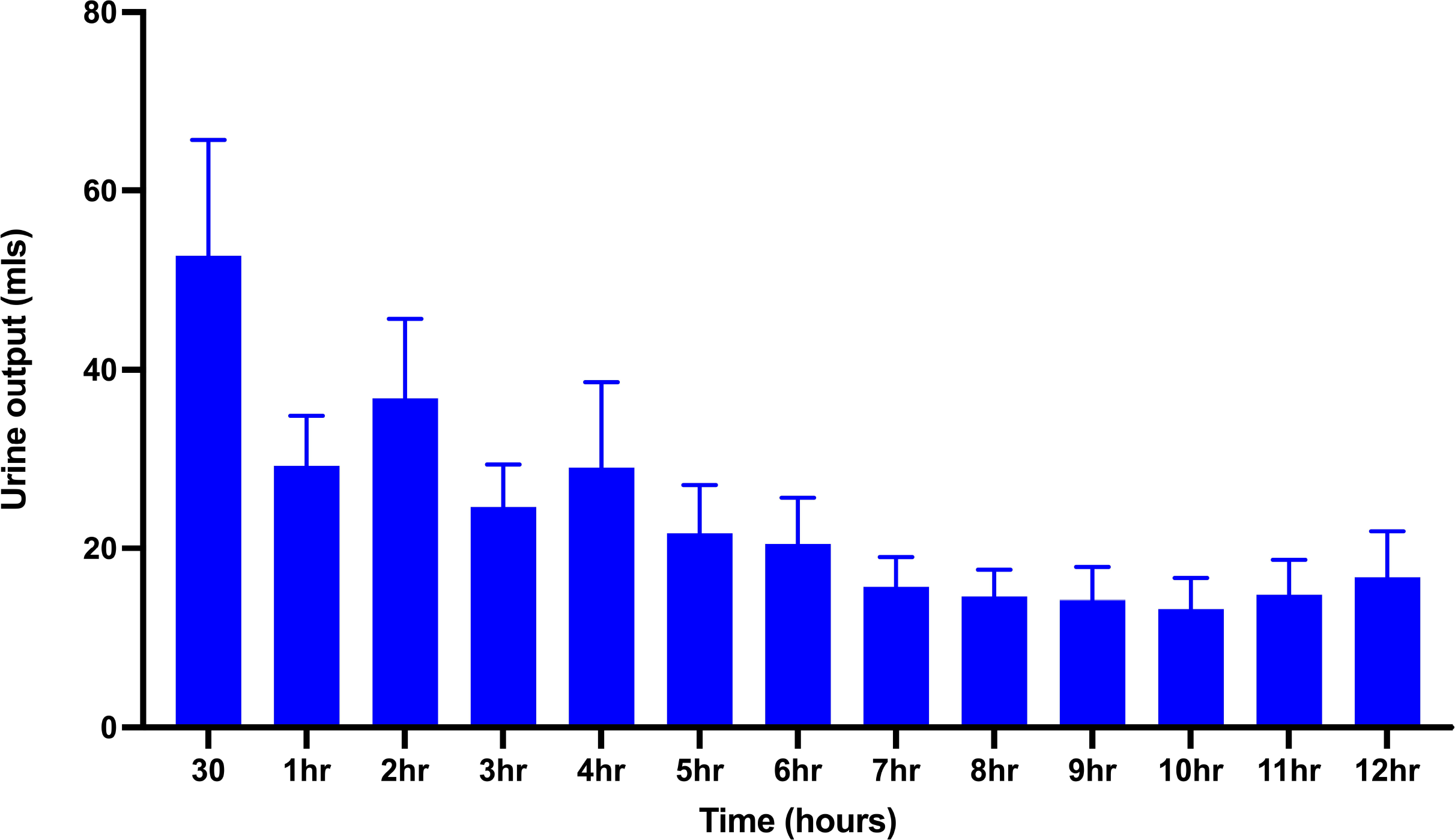
Kidney function is restored. All kidneys began producing urine immediately as they were connected to the NMP circuit, and this was maintained throughout the experiment.

### Clinical Assessment Score

All kidneys presented with a globally pink appearance which was maintained throughout. Following 1 hour of stabilisation, RBF was consistently above 50ml/hr/100g and total UO was >43mls. This provides all kidneys with a combined total score of 1 where they would be deemed acceptable for transplantation.

### Histological results

To assess tissue integrity, all biopsies were H&E stained and analysed by a veterinary histopathologist. Kidney samples were graded according to the Remuzzi scoring system. There was no evidence of glomerular global sclerosis, tubular atrophy, interstitial fibrosis or vascular narrowing in any of the renal samples, giving all kidneys a final Remuzzi score of 0 and indicating suitability for single transplant (Figure 5A and 5B). Finally, gram staining revealed no bacteria present in any renal samples.

**Figure 5.**
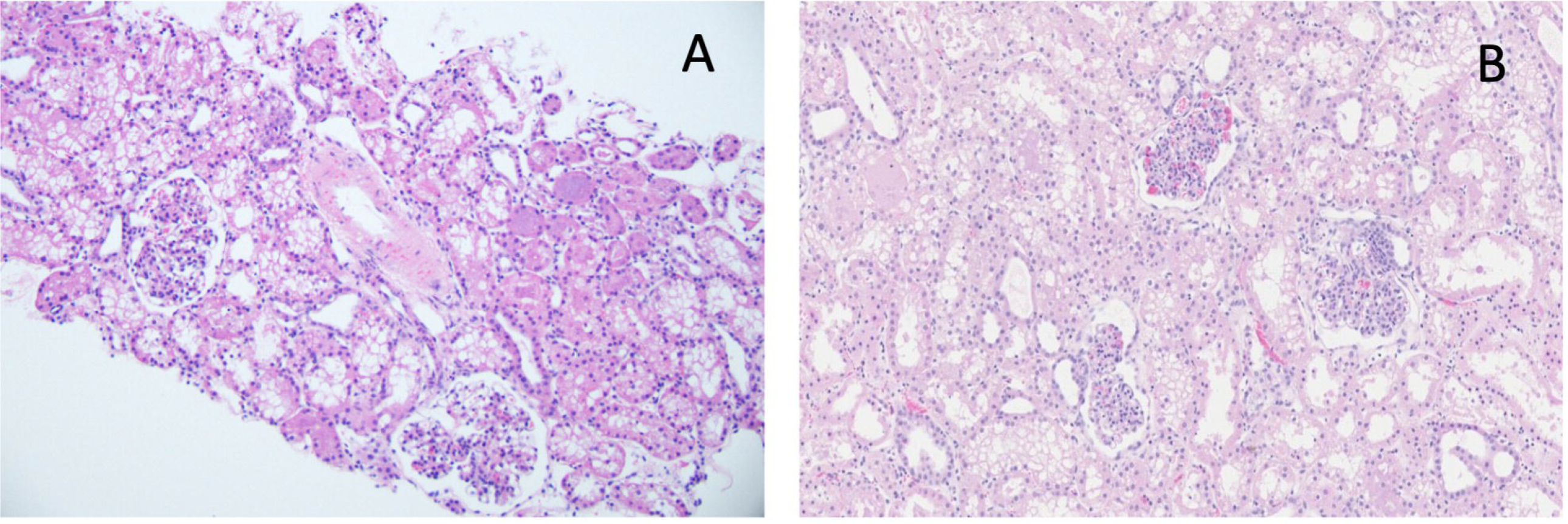
Representative H&E histology images. A) 20x magnification of a core needle biopsy. B) 15x magnification of a wedge biopsy.

### Weight Change

The average weight of the kidneys before they were perfused was 331.4g ± 35.85. Following 12 hours of perfusion, this was 397.2g ± 50.17. Overall, the percentage weight change was 19.41% ± 10.30, most likely due to the increased blood-perfusate present in the kidney (Figure 6).

**Figure 6.**
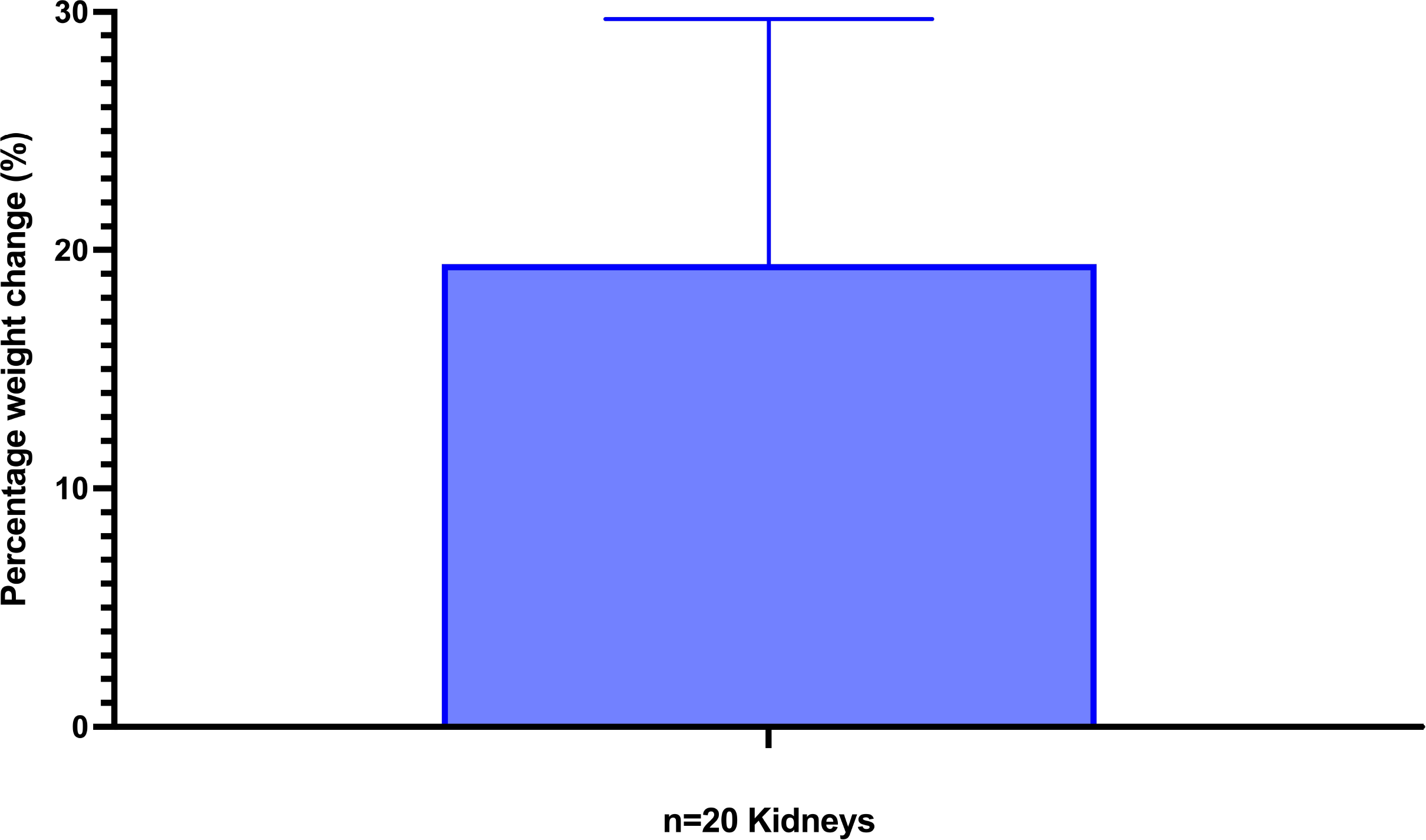
Percentage weight change between pre and post perfusion. The percentage weight change for all kidneys was 19.41% ± 10.30.

## 4 Discussion

NMP offers a novel platform for the preservation, optimisation and evaluation of donor kidneys prior to transplantation. Currently, a validated protocol does not exist beyond 3 hours for clinical purposes (4,7,8). We have developed a NMP protocol that can maintain a donor kidney for 12 hours. In all kidneys, successful perfusion was achieved with stable haemodynamics and blood-perfusate biochemistry. RBF and IRR improved continuously throughout, without fluctuations in arterial pressure, indicating improved vascular tone and endothelial integrity of the renal vasculature. Electrolyte balance and haematocrit were maintained at physiological levels, along with sustained, and physiological urine output. Previous studies of prolonged kidney NMP, without urine recirculation, have reported disruption of electrolytes, particularly elevated sodium levels (9,10). In the kidney, hypernatremia is vasoconstrictive, causing a reduction in renal arterial blood flow and increasing pressure (11); which in turn impacts glomerular filtration rate. Therefore, the maintenance of sodium levels within physiological range in this protocol are of particular importance.

Metabolic function was restored, including renal gluconeogenesis. A continuous reduction in blood-perfusate lactate was observed with time. Lactate clearance is a key metabolic process, with high lactate levels indicative of anaerobic metabolism and hypoperfusion. Indeed, a lactate level of less than 5mmol/L at the end of perfusion has been proposed as a key criterion for organ transplantation (3). As such, the maintenance of lactate levels below 2mmol/L in this protocol demonstrates sustained aerobic metabolism, tissue health and suitability for transplantation.

Pyruvate produced during gluconeogenesis is also converted to acetyl-CoA for the citric acid (TCA) cycle, forming bicarbonate as a by-product. Continuous generation of bicarbonate was recorded, suggesting active pyruvate to bicarbonate conversion. This is supported by the significant negative correlation between lactate and bicarbonate concentrations within the perfusate. Bicarbonate recycling from lactate is required to maintain acid/base balance and buffer pH. The slightly alkalotic pH recorded at the end of perfusion can thus be attributed to sustained gluconeogenesis and pyruvate to bicarbonate conversion. Sodium bicarbonate was only supplemented as required during stabilisation within the first 30 minutes of perfusion. Following this, kidneys maintained acid-base balance via biochemical processing and no further supplementation was required.

As cellular metabolism is restored under the described protocol, NMP provides the opportunity to assess graft function and viability prior to transplantation, potentially in the context of improving physiological status. All kidneys achieved a score of 1 on the established clinical grading system following 12 hours of perfusion, suggesting high quality preservation and a status equating to suitability of a perfused human kidney for transplantation (3). In addition to the assessment criteria proposed by Hosgood et al. macroscopic dissection of kidneys at the end of perfusion revealed active calyceal pacing and no obvious signs of damage to the cortex or medulla, together indicating optimal preservation and tissue integrity. This is further evidenced by the H&E histology of kidney biopsies confirming excellent preservation of tissue health and absence of injury after 12 hours NMP in both groups. A wider qualitative assessment made on the general state of tissue integrity, did not find any evidence of acute injury. Other studies of kidney NMP have found histopathological damage in the form of acute ischemic changes, tubular dilation, tubular debris, vacuolation and infiltration (12,13). It cannot be determined if histology improved throughout perfusion as baseline biopsies were not taken.

The ability to safely preserve a donor kidney for 15.5hrs (based on 3.5hrs cold storage and 12hrs perfusion has the capacity to significantly improve renal transplantation logistics. Transplantation services are severely limited by resource issues, intensity of night work, demand on shared theatres and recipient delays. Extending preservation via SCS is not feasible, as cold ischaemia is directly related to the severity of reperfusion injury, with each hour of SCS associated with an increased risk of DGF (2,14). In practise, this means many kidney transplant units are currently forced to perform the majority of often complex transplant surgeries during the night, which has many disadvantages. Extending non-injurious preservation via NMP, beyond the currently applied protocols could provide the opportunity to defer all such cases to the daytime with all the safety and health-and wellbeing benefits that come with that. This potential “electivisation” of renal transplantation has major potential benefits beyond patient safety including recruitment/retention and longevity of transplant surgeons, cost improvements through reduced out-of-hours cover requirement and third-party patient benefit from freeing emergency theatre space.

Whilst we have developed a 12-hour perfusion protocol with maintenance of key physiological parameters, there remain some key limitations of this study. The study used healthy pig kidneys, whilst clinical kidney transplantation often involves the use of kidneys from older patients with significant chronic comorbidity and the established acute effects of a terminal illness. How such kidneys will respond to prolonged optimised NMP will require further study.

## Summary

In this study we confirm the feasibility for prolonged perfusion with NMP to 12 hours, maintaining donor kidneys in optimal health. Multiple aspects of renal physiology were confirmed to be maintained in a near normal state including renal haemodynamics, blood-perfusate biochemistry, urine production and macroscopic appearance in 20 kidneys. We now intend to evaluate the protocol following 12 hours SCS, to extend NMP to 24 hours, and to determine post-transplant outcomes compared to SCS alone. Extending the timeframe that donor kidneys can be preserved using this technology has multiple benefits including the ability to schedule transplants, treat kidneys and reduce ischaemic injury.

## Author Contributions

JPS – *Significantly contributed to the conception and design of the study, the acquisition, analysis and interpretation of all data; (ii) drafting the article and revising it critically for important intellectual content; and (iii) has approved the final version for publication*

WRC - *Significantly contributed to the acquisition, analysis and interpretation of data; (ii) drafting the article and revising it critically for important intellectual content; and (iii) has approved the final version for publication*

CJTB - *Significantly contributed to the acquisition, analysis and interpretation of data; (ii) revising the article critically for important intellectual content; and (iii) has approved the final version for publication*

AFS- *Significantly contributed to the acquisition, analysis and interpretation of data; (ii) revising the article critically for important intellectual content; and (iii) has approved the final version for publication*

ERA- *Significantly contributed to the acquisition, analysis and interpretation of surgical data and (ii) has approved the final version for publication*

MC*- Significantly contributed to the acquisition, analysis and interpretation of surgical data and (ii) has approved the final version for publication*

TRE - *Significantly contributed to the acquisition, analysis and interpretation of surgical data and (ii) has approved the final version for publication*

JDP *- Significantly contributed to the acquisition, analysis and interpretation of surgical data*

KA - *Significantly contributed to the conception and design of the study, the acquisition, analysis and interpretation of data; (ii) drafting the article and revising it critically for important intellectual content; and (iii) has approved the final version for publication*

JEF- *Significantly contributed to the conception and design of the study, the acquisition, analysis and interpretation of data; (ii) drafting the article and revising it critically for important intellectual content; and (iii) has approved the final version for publication*

## Data Availability Statement

All datasets generated for this study are including in this article and are stored on a secure electronic laboratory notebook.

## References

1. NHSBT. Annual Report on Kidney Transplantation. 2022.

2. Debout A, Foucher Y, Trébern-Launay K, Legendre C, Kreis H, Mourad G, et al. Each additional hour of cold ischemia time significantly increases the risk of graft failure and mortality following renal transplantation. Kidney Int. 2015 Feb 3;87(2):343–9.

3. Hosgood SA, Barlow AD, Hunter JP, Nicholson ML. Ex vivo normothermic perfusion for quality assessment of marginal donor kidney transplants. British Journal of Surgery. 2015 Sep 9;102(11):1433–40.

4. Nicholson ML, Hosgood SA. Renal transplantation after ex vivo normothermic perfusion: the first clinical study. Am J Transplant. 2013 May;13(5):1246–52.

5. Stone JP, Ball AL, Critchley WR, Major T, Edge RJ, Amin K, et al. Ex Vivo Normothermic Perfusion Induces Donor-Derived Leukocyte Mobilization and Removal Prior to Renal Transplantation. Kidney Int Rep. 2016;1(4):230–9.

6. Remuzzi G, Grinyò J, Ruggenenti P, Beatini M, Cole EH, Milford EL, et al. Early Experience with Dual Kidney Transplantation in Adults Using Expanded Donor Criteria. Journal of the American Society of Nephrology. 1999 Dec;10(12):2591–8.

7. Hosgood SA, Saeb-Parsy K, Wilson C, Callaghan C, Collett D, Nicholson ML. Protocol of a randomised controlled, open-label trial of ex vivo normothermic perfusion versus static cold storage in donation after circulatory death renal transplantation. BMJ Open. 2017 Jan 1;7(1):e012237.

8. Rijkse E, De Jonge J, Kimenai HJAN, Hoogduijn MJ, De Bruin RWF, Van Den Hoogen MWF, et al. Safety and feasibility of 2 h of normothermic machine perfusion of donor kidneys in the Eurotransplant Senior Program. BJS Open. 2021 Jan 1;5(1):zraa024.

9. Weissenbacher A, Bogensperger C, Oberhuber R, Meszaros A, Gasteiger S, Ulmer H, et al. Perfusate Enzymes and Platelets Indicate Early Allograft Dysfunction After Transplantation of Normothermically Preserved Livers. Transplantation. 2022 Apr 1;106(4):792–805.

10. Weissenbacher A, Lo Faro L, Boubriak O, Soares MF, Roberts IS, Hunter JP, et al. Twentyfour–hour normothermic perfusion of discarded human kidneys with urine recirculation. American Journal of Transplantation. 2019 Jan 1;19(1):178–92.

11. Osswald H, Gleiter C. [Hypernatremia and kidney function]. Zentralbl Chir. 1993;118(5):267–72.

12. Hosgood S, Harper S, Kay M, Bagul A, Waller H, Nicholson ML. Effects of Arterial Pressure in an Experimental Isolated Haemoperfused Porcine Kidney Preservation System. British Journal of Surgery. 2006 Jun 14;93(7):879–84.

13. Hosgood SA, Barlow AD, Yates PJ, Snoeijs MGJ, Van Heurn ELW, Nicholson ML. A Pilot Study Assessing the Feasibility of a Short Period of Normothermic Preservation in an Experimental Model of Non Heart Beating Donor Kidneys. Journal of Surgical Research. 2011 Nov 1;171(1):283–90.

14. Patel K, Nath J, Hodson J, Inston N, Ready A. Outcomes of donation after circulatory death kidneys undergoing hypothermic machine perfusion following static cold storage: A UK population-based cohort study. American Journal of Transplantation. 2018 Jun 1;18(6):1408–14.

